# Investigating the translational value of Periprosthetic Joint Infection (PJI) models to determine the risk and severity of Staphylococcal biofilms

**DOI:** 10.1101/2024.04.29.591689

**Authors:** Amita Sekar, Yingfang Fan, Peyton Tierney, Madeline McCanne, Parker Jones, Fawaz Malick, Devika Kannambadi, Keith K Wannomae, Nicoletta Inverardi, Orhun Muratoglu, Ebru Oral

**Author notes:** These authors contributed equally (Amita Sekar, Yingfang Fan).

## Abstract

With the advent of antibiotic-eluting polymeric materials for targeting recalcitrant infections, using preclinical models to study biofilm is crucial for improving the treatment efficacy in periprosthetic joint infections. The stratification of risk and severity of infections is needed to develop an effective clinical dosing framework with better outcomes. Here, using in-vivo and in-vitro implant-associated infection models, we demonstrate that methicillin-sensitive and resistant *Staphylococcus aureus* (MSSA and MRSA) have model-dependent distinct implant and peri-implant tissue colonization patterns. The maturity of biofilms and the location (implant vs tissue) were found to influence the antibiotic susceptibility evolution profiles of MSSA and MRSA and the models could capture the differing host-microbe interactions in vivo. Gene expression studies revealed the molecular heterogeneity of colonizing bacterial populations. The comparison and stratification of the risk and severity of infection across different preclinical models provided in this study can guide clinical dosing to effectively prevent or treat PJI.

## Introduction

Periprosthetic joint infections (PJIs) represent a formidable challenge to the success of total joint replacements. These infections, characterized by microbial colonization on the implant materials and the surrounding tissue, not only compromise the intended function of the implants but also significantly reduce the quality of life of the patients [1–3]. In addition, they are increasingly hard to treat with recurrent infections causing significantly increased morbidity and mortality [4].

Currently, a generalized approach is used in treating suspected PJI, where one of several avenues of treatment is utilized [5,6]. While systemic antibiotics are the main tool in addressing bacterial infections, the implants can be retained or replaced in a one- or two-stage revision [7]. Locally, the elution of aminoglycosides together with vancomycin from antibiotic-eluting bone cement is used to support the role of systemic antibiotics [8]. It is believed that the high concentrations achieved by local administration/elution of drugs will lead to higher efficacy in addressing joint infections while reducing systemic side effects of antibiotics such as nephrotoxicity. However, there is little conclusive information on the precise effects of local antibiotic administration and their efficacy in preventing/treating PJI and there is no specific dosing guidance for local antibiotics [9–13]. Commonly used dosing for prophylaxis may lead to the worsening of outcomes [14]. Thus, there is a great need to determine the dosing requirements for antibiotics to prevent and treat PJI locally.

Methicillin-susceptible (MSSA) and methicillin-resistant (MRSA) *S. aureus* have been established as the most significant causative organisms of periprosthetic joint infections [15,16]. These bacterial strains adhere to inanimate surfaces and the surrounding tissue using specific cell surface proteins and adhesins such as elastin-binding proteins[17]. Further aggregation triggers bacterial regulatory pathways promoting the production of extracellular polymeric substances such as polysaccharide intercellular adhesion (PIA) regulated by *ica* operon) that establishes robust biofilms[18]. The microbial communities in biofilm status undergo significant molecular, physiological, and morphological changes which provide bacteria with resilience against host defenses and conventional antibiotics such as gentamicin[19,20]. These structured microbial communities, with their cell wall modifications and an intricate architecture comprising live and dead bacterial aggregates together with host cell components embedded in a complex polymeric matrix, create diffusion barriers leading to a high tolerance to antibiotics and host factors[21]. They also serve as a reservoir for maintaining the hardy bacterial populations that can persist in the presence of antibiotics and infiltrate deep tissue spaces[22,23]. Heterogenous subpopulations of bacteria in the biofilm present varying phenotypic and genotypic profiles within the same biomass. These subpopulations differentially specialize in pathogenesis, drug resistance, and evading immune responses[24,25]. Due to the reasons mentioned above, antibiotic-based treatments are often not effective against these diverse populations and physical removal (radical debridement) is necessary for definitive treatment. Despite debridement, antibiotic lavage, replacement of implant components, and local elution from antibiotic-eluting bone cement, failure rates are high in PJI[26–28].

While the significance of biofilms in periprosthetic joint infections is acknowledged[29,30], there remains a substantial knowledge gap concerning the infection dynamics, phenotypic and genotypic heterogeneity, and the time-dependent properties of antibiotic susceptibility. In-vitro models may not capture the biofilm properties in response to a realistic environment, despite providing a straightforward approach to studying biofilms. More recently, studies have attempted to characterize and correlate biofilm formation and antibiotic susceptibility of clinical isolates to outcomes of PJI[31–33]. However, the characterization of ex vivo-grown bacterial cultures is also limited in capturing the susceptibility dynamics in vivo. There is also little information on emerging heterogenous populations based on colonizing location (implant surface vs tissue), vascularization, and biofilm physiology. There is a need for a better understanding of the infecting organism together with their innate and adaptive behavior within a PJI setting to determine the risk and severity of the infection. This crucial knowledge can aid in the design and development of multi-faceted drug-eluting materials and treatment algorithms.

Our long-term goal is to design antibiotic-eluting polymeric materials that can be more efficiently used locally to prevent and treat PJI. As material advancements and preclinical research enhance our understanding and prevention of periprosthetic joint infections (PJI), notable discrepancies are exposed between promising preclinical findings and clinical testing outcomes. One factor contributing to the lack of predictability of the clinical outcomes associated with PJI treatment is the variability in the bacterial strain and the timing of treatment based on clinical symptoms[34,35]. In this study, we developed an in-vitro implant material infection model, a subcutaneous implantation and infection model, and a periprosthetic joint infection model in the rat, modeling *S. aureus* infections with varying risk and severity. To capture the range of therapeutic dosing dependent on bacterial evolution, we proposed to use two *S. aureus* bacterial strains, with and without inherent resistance to the aminoglycoside gentamicin. These preclinical models were designed to monitor bacterial dynamics and bacterial resistance evolution to gentamicin and to understand the molecular events within biofilms contributing to resistance and persistence. We hypothesized that we could capture a ‘therapeutic window’, providing a guideline for local dosing to prevent or treat PJI for ‘low-risk’ and ‘high-risk’ infections clinically. A secondary goal was to compare the bacterial dynamics and resistance evolution in vitro and in vivo in preclinical models to enhance the translational value of antibacterial testing of antibiotic-eluting polymeric materials.

## RESULTS

### MSSA and MRSA demonstrate distinct colonization patterns in in-vivo and in-vitro models

The biofilm localization and growth dynamics of MSSA and MRSA were determined on both the implant material and the peri-implant tissue from the subcutaneous and joint infection models. In the subcutaneous model, the viability of implant-adherent bacteria recovered was consistently 10^3^ CFU/mL for 21 days. There were≥2log more viable bacteria recovered from peri-implant tissue samples, with the viable load being highest at POD 1 and 3 (10^8^ CFU/mL), which was subsequently reduced by POD 21 (10^5^ CFU/mL) (Figure 1A). SEM observations confirmed poor bacterial presence/viability on the surface of the implanted plates. MRSA demonstrated more bacterial aggregates and biofilm matrix structures on SS plates when compared to MSSA (Figure 2A).

**Figure 1.**
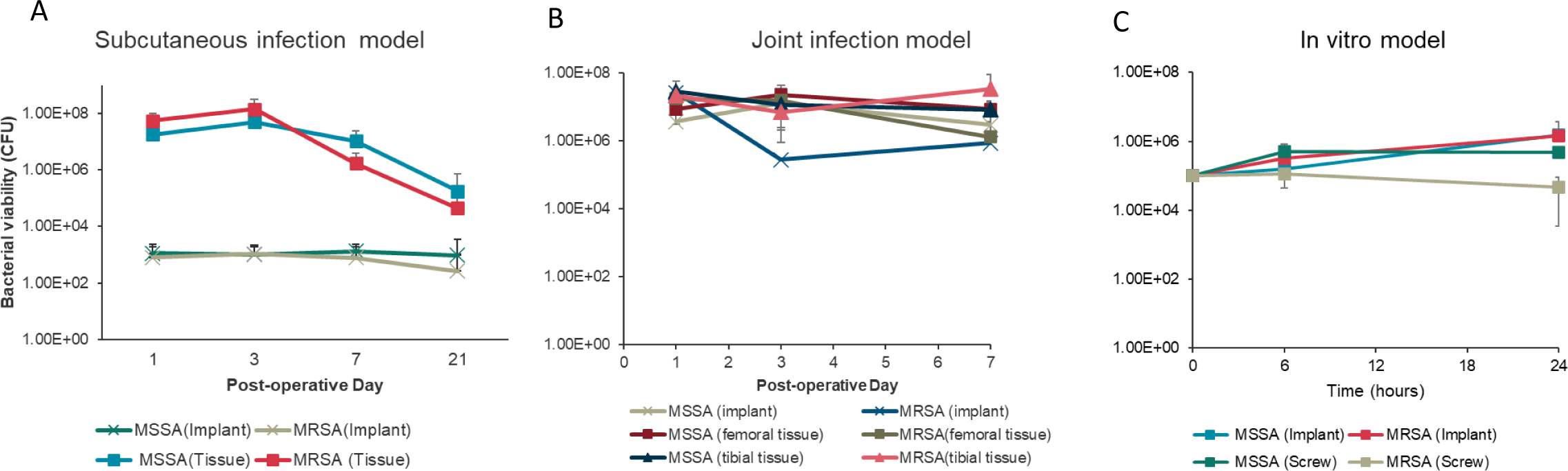
Biofilm growth dynamics of MSSA and MRSA across in vivo and in vitro infection models. (A) MSSA and MRSA count from stainless-steel implant and peri-implant tissue harvested at POD 1, 3, 7, and 21. (B) MSSA and MRSA count from stainless-steel screw implant and peri-implant femoral and tibial tissue harvested at POD 1, 3, and 7. (C) Adherent MSSA and MRSA bacteria count from in-vitro stainless-steel plate and screw culture at 6 and 24 hours. Bacterial viability data from tissue samples were calculated by normalizing to the respective weight of tissue retrieved. Error bars represent the standard deviation (n=2 for implant, n=4 for tissue culture in (A), n=1 for screw implant, n=3 for tissue culture in (B) and n=3 in (C)). n indicates the number of samples.

**Figure 2:**
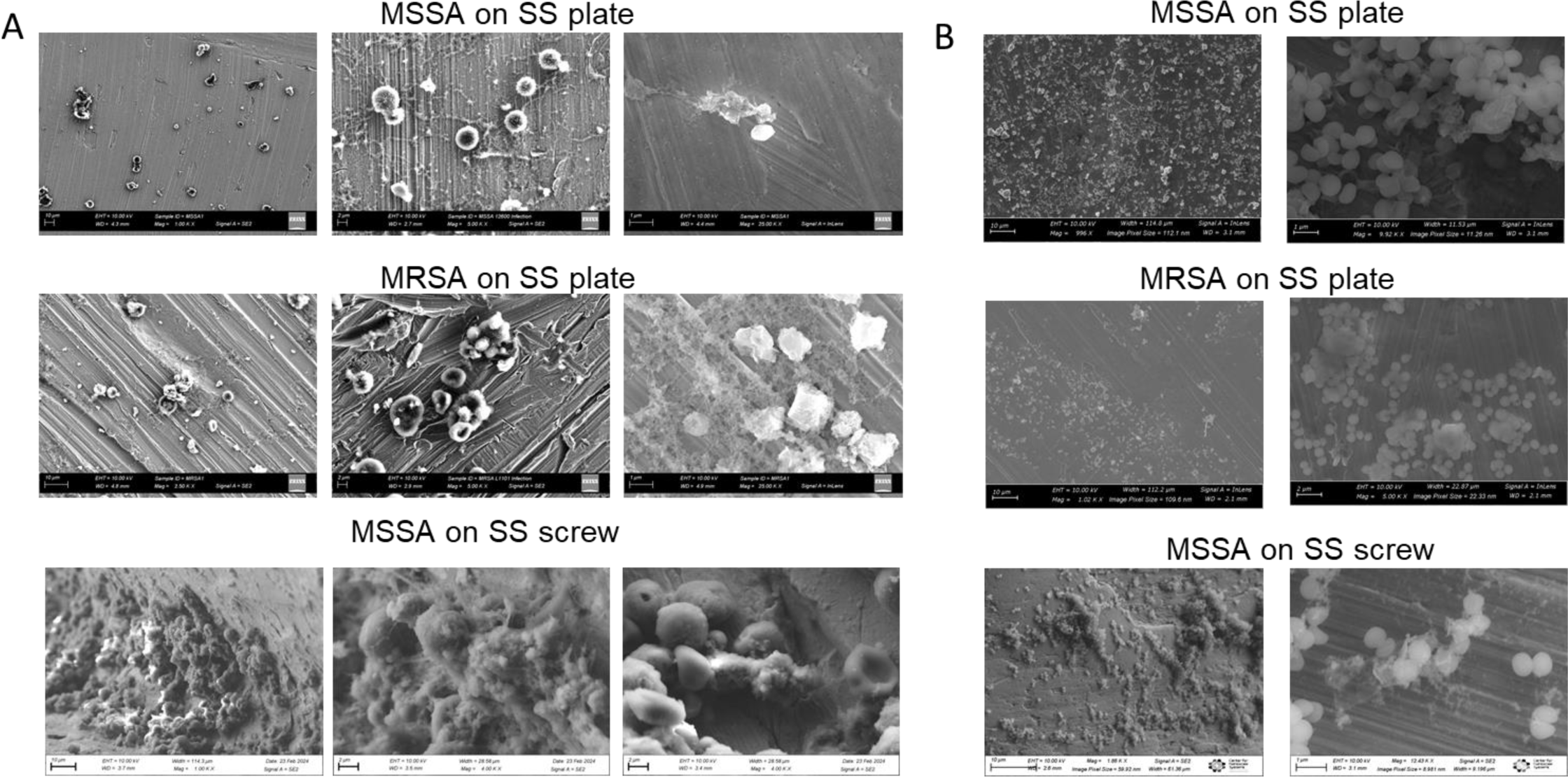
*S. aureus* adhesion on implant materials retrieved from in vivo and in vitro models. Scanning electron micrographs of A) MSSA and MRSA adhered to implanted stainless-steel materials retrieved on POD3 from in vivo infection models and B) Established (24 hours) biofilms of MSSA and MRSA adhered to stainless-steel materials in vitro. Scale bar = 10, 2, and 1μm for panel A and 10 and 2 μm for panel B respectively. Representative images of n=2 implants from subcutaneous and joint infection model; n=3 for in vitro model

High MSSA and MRSA bacterial viability (>5×10^5^ CFU/mL) was observed in the joint infection model on both screw implant and peri-implant tissue (Figure 1B). The MRSA on the screw implant showed lower viability (∼2 log) on POD 3 compared to MSSA but the bacterial load was similar for both strains at POD 7. The bacterial viability of MSSA recovered from the peri-implant femoral and tibial tissues did not show any differences. However, the viable MRSA recovered from the peri-implant (tibia) tissue was consistent over time (∼10^7^ CFU/mL) when compared to that of peri-implant femoral tissue which steadily decreased over the period of 7 days (from >5 ×10^7^ to 1×10^6^ CFU/mL). SEM confirmed increased bacterial adherence to the screw implant surface. Significant adhesion and biofilm formation comprising varying cell morphologies and dense matrix components were observed for MSSA on the implanted surface (Figure 2B).

In the in-vitro model, the viable bacteria recovered from MSSA-adhered SS plates and screws increased from ∼10^5^ to > 10^6^ CFU in 24 hours (Figure 1C). SEM observations showed significant bacterial attachment on implant materials (SS plates and screws) which correlated with the bacterial viability data (Figure 2B).

### Biofilm formation and dynamics influence bacterial susceptibility to gentamicin

The susceptibility of tissue-colonized and implant-adherent MSSA and MRSA to the antibiotic gentamicin was determined longitudinally. In the subcutaneous model, the implant-adhered MRSA demonstrated antibiotic resistance as early as POD 1 (200 µg/mL). At POD 3 and until POD 21, the MBEC was found to be above the maximum concentration of gentamicin tested in this study (>500 µg/mL). Implant-adhered MSSA acquired resistance towards gentamicin by POD 7 (100 µg/mL), which remained the same until POD 21 (Figure 3A, Table 1). The tissue-colonized MRSA and MSSA exhibited the highest resistance (>500 and 200 µg/mL, respectively) on POD 1 and 3, which was decreased by POD 21 (200 µg/mL and no growth, respectively) (Figure 3A, Table 1).

**Figure 3:**
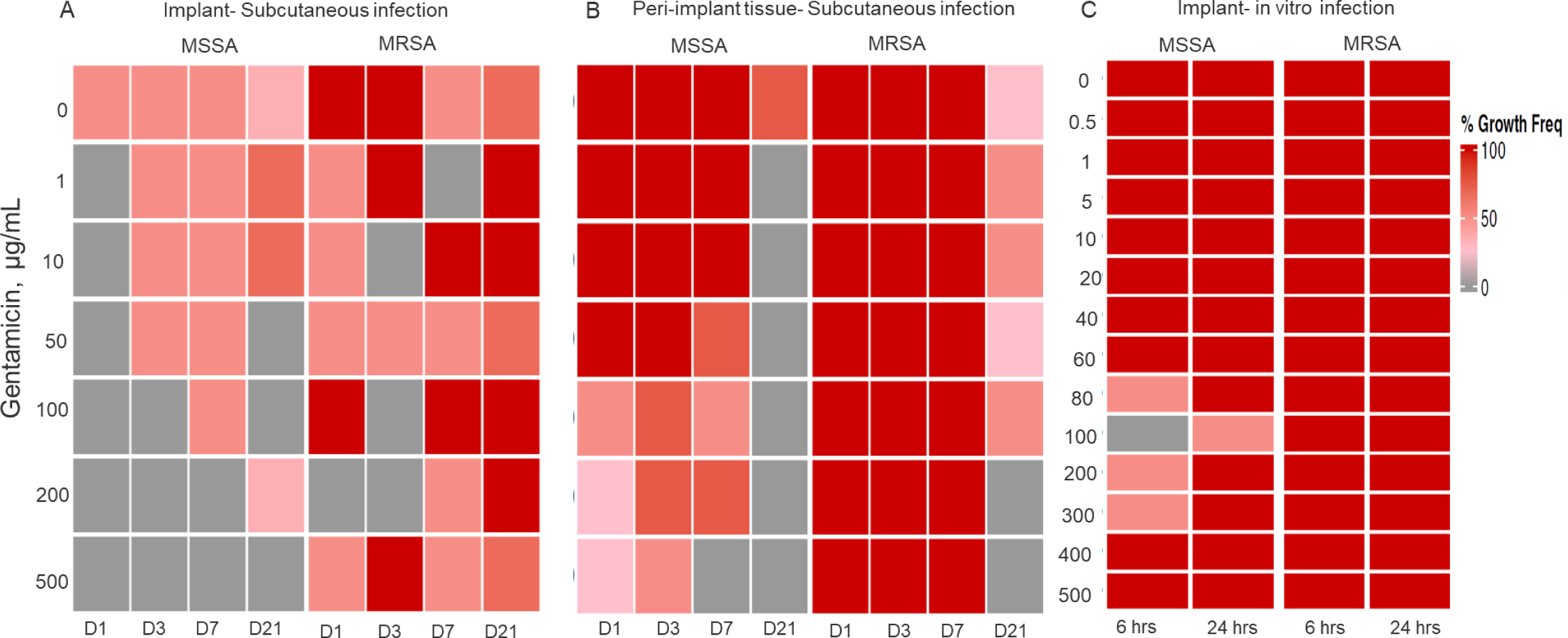
Evolution of gentamicin susceptibility for MSSA and MRSA. Heat maps indicating the %growth frequency observed after 24 hours of indicated gentamicin concentration exposure for A) MSSA and MRSA adhered to the subcutaneously implanted stainless steel material retrieved on POD1,3,7and 21 (POD1,3,7 n=2; POD21 n=3) B) MSSA and MRSA colonizing peri-implant tissue retrieved on POD1,3,7 and 21 (POD1,3,7 n=4; POD21 n=2) C) Nascent (6 hours) and established (24 hours) biofilms of MSSA and MRSA adhered to stainless steel material in vitro (n=3). The susceptibility profiles of gentamicin for implant and tissue samples across the different models are presented in Tables 1-3.

**Table 1:**
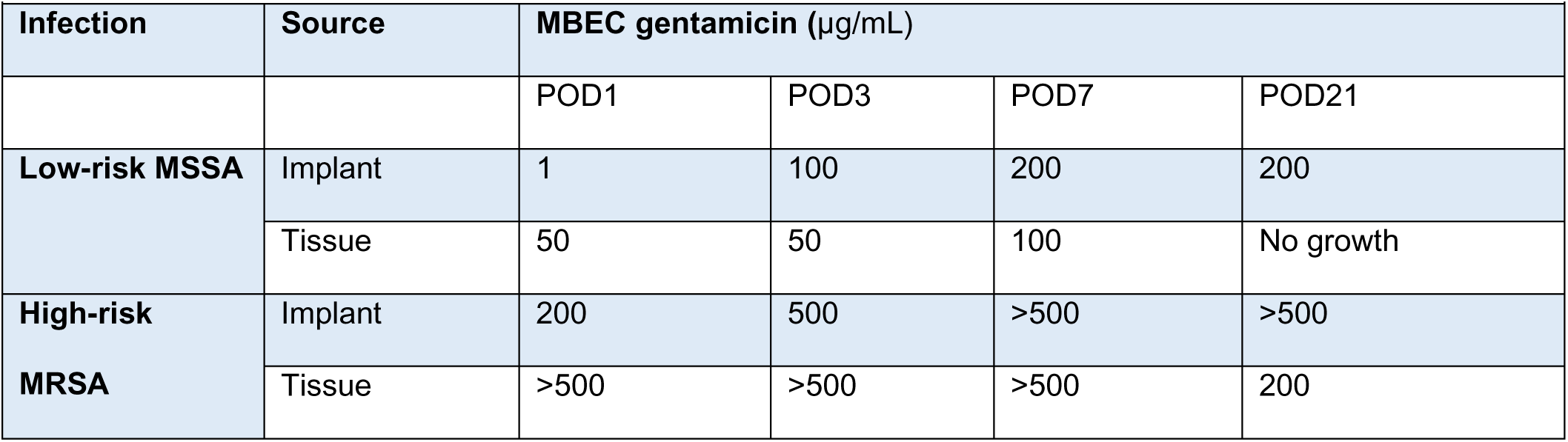
Subcutaneous infection model.

In the joint infection model, the implant-adhered MRSA demonstrated very high antibiotic resistance as early as POD 1 (>500 µg/mL) and stayed highly resistant until POD 21. In contrast, the implant adhered MSSA demonstrated a comparatively lower level of acquired resistance (≥10 µg/mL) at POD 1, 3, and 7 (Figure 3B). For the tissue colonized MSSA, the acquired gentamicin resistance was moderate and consistent over 7 days of infection (50 µg/mL). The tissue colonized MRSA showed the highest resistance on POD 1 and 3 and a subsequent reduction of MBEC to 300 µg/mL was observed by POD 7 (Table 2).

**Table 2:** Joint infection model.

| Table 2: Joint infection model |  |  |  |  |
| --- | --- | --- | --- | --- |
| Infection | Source | MBEC gentamicin (µg/mL) |  |  |
|  |  | POD1 | POD3 | POD7 |
| Low-risk MSSA | Implant | >10 | >10 | 10 |
|  | Tissue | 50 | 50 | 50 |
| High-risk MRSA | Implant | >500 | >500 | >500 |
|  | Tissue | >500 | >500 | 300 |

For the in-vitro SS plate model, both implant-adhered MSSA and MRSA demonstrated increased resistance due to biofilm formation within 6 hours (100 and >500 µg/mL, respectively). The acquired resistance of MSSA was further increased to >500 µg/mL within 24 hours, whereas MRSA stayed highly resistant throughout the study period (Figure 3C, Table 3). For the in-vitro SS screw model, both 6 hours and 24 hours-grown biofilms of MSSA exposed to gentamicin were eradicated effectively with concentrations close to its planktonic MIC values (5 µg/mL). The 6-hour and 24-hour biofilms of MRSA grown on SS screws exhibited inherent resistance to gentamicin (3log reduction only at 500 µg/mL and no reduction until 500 µg/mL, respectively).

**Table 3:** in vitro model.

| Table 3: in vitro model |  |  |  |
| --- | --- | --- | --- |
| Infection | Source | MBECgentamicin |  |
|  |  | Nascent (6hr) | Established (24hr) |
| Low-risk MSSA | Implant | 100 | >500 |
|  | Screw | 5 | 5 |
| High-risk MRSA | Implant | >500 | >500 |
|  | Screw | 500 | >500 |

### Molecular responses of bacterial biofilms to their environment are strain-dependent

The gene expression of MSSA and MRSA colonizing the implant materials (in vivo and in vitro) and the peri-implant tissue were performed to reveal their molecular status. In the subcutaneous model, the *vraR* gene expression for tissue colonized MSSA was elevated (>1.5log_10_) for the length of the study. For MRSA, the *vraR* expression was not altered until POD 7, when the expression was deregulated but the expression was significantly increased on POD 21 (>1.5log_10_). The *icaA* gene expression of MSSA was found to be largely unaltered until POD 21 where it was somewhat increased (∼1log_10_). For MRSA, the *icaA* gene was upregulated significantly at POD 7 (>1 log_10_) and remained elevated on POD 21. The *icaD* gene expression in both MSSA and MRSA was largely unaltered over the entire study period (< 0.5 log_10_). The *ebpS* expression demonstrated a consistent increase for both MSSA and MRSA from POD 1 to POD 21 (>0.5 log_10_) (Figure 4A).

**Figure 4:**
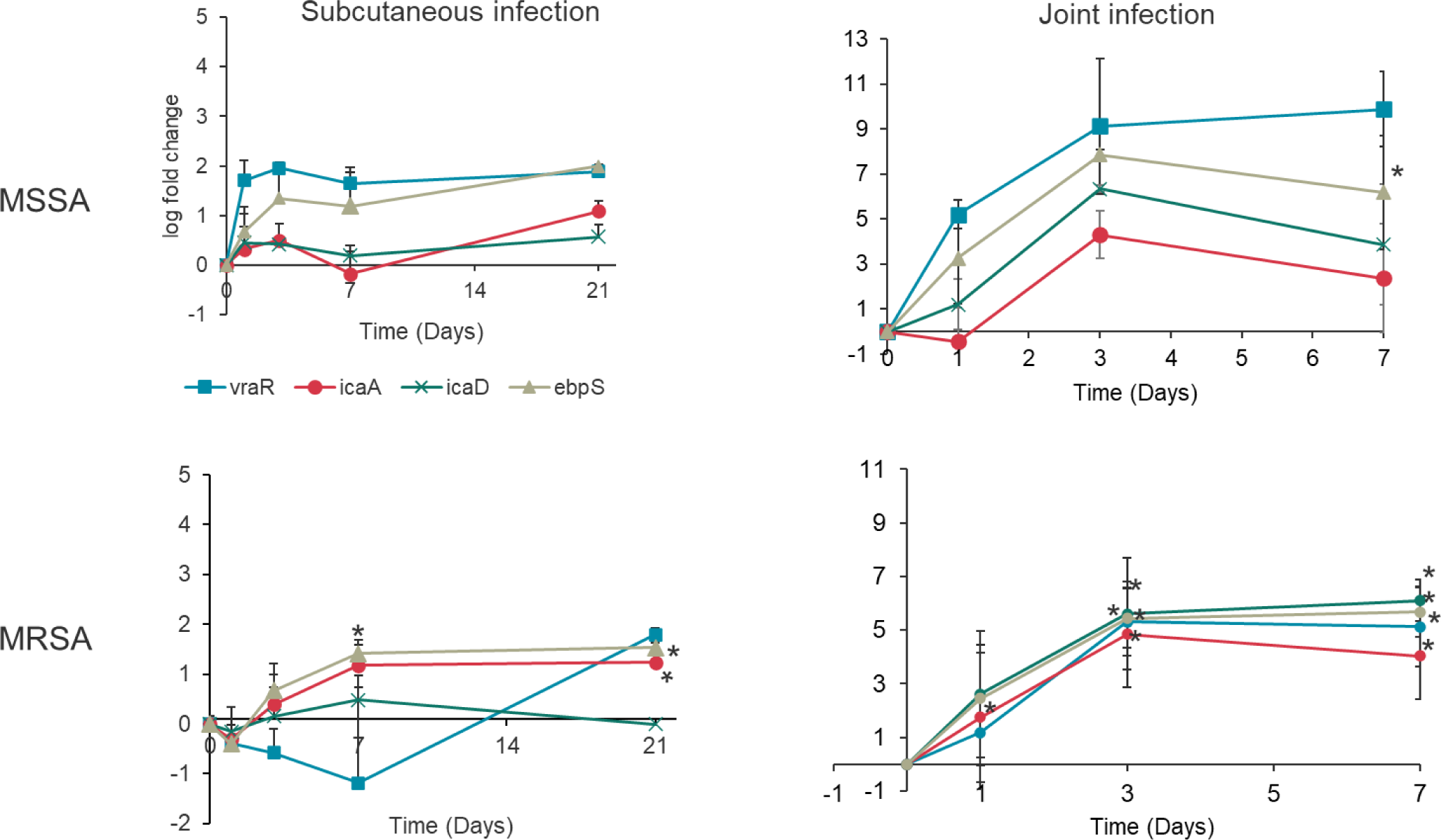
Regulation of stress responses, adhesion, and biofilm formation in tissue colonizing MSSA and MRSA in vivo. Gene expression analysis of A) MSSA and MRSA colonizing the peri-implant tissue retrieved on POD1,3,7 and 21 from rats subjected to subcutaneous infection and POD1,3,7 from rats subjected to joint infection. B) MSSA and MRSA biofilms adhered to the implant materials retrieved on POD 3 from in vivo models and from in vitro models at time points indicated. The relative gene expression was further normalized to the expression profile of planktonic *S. aureus* suspension. Error bars represent standard deviation (Subcutaneous model; n=3 (POD 1,3,7) n=10 and 11 for POD21 MSSA and MRSA respectively, Joint model; n=3 (POD1,3,7) *Indicates p value <0.05

In the joint infection model, the *vraR* expression for MSSA was found to be significantly upregulated (>2log_10_) and the expression levels were highest on POD7 (>3.5 log_10_). For MRSA, *vraR* expression was slightly altered on POD 3 and 7 (>0.5 log_10_). The *icaA* and *icaD* genes for MSSA were significantly upregulated on POD 3 and 7 (>1 log_10_). For MRSA, the *icaA* expression was somewhat upregulated (∼0.5 log_10_) and the *icaD* expression was significantly upregulated until POD 7 (>1 log_10_). The *ebpS* gene expression was significantly increased for MSSA with the highest expression demonstrated on POD 3 and 7 (>2 log_10_). For MRSA, the *ebpS* gene expression was increased (∼1 log_10_) with overall subdued gene expression in comparison to MSSA (Figure 4B).

In the in-vitro model, the implant-adhered MSSA demonstrated downregulation of *vraR* and *ebpS* genes (−1.5 log_10_) in nascent biofilms (6 hours) and the expression remained unaltered in established biofilms (24 hours). For MRSA, both genes were significantly downregulated in nascent biofilms and the expression remained the same in established biofilms (>-1.5 log_10_). The *icaA* and *icaD* expression was found to be drastically deregulated in both MSSA and MRSA harvested from nascent and established biofilms (>-3 log_10_) (Figure 5). In the in-vivo models (subcutaneous and joint infection) the biofilm RNA extracted from implant materials demonstrated strain-dependent expression of 16srRNA indicating the presence of bacteria on the surfaces. The expression profile for the remaining genes was not observed. (Table 5, Supplementary File 1).

**Figure 5:**
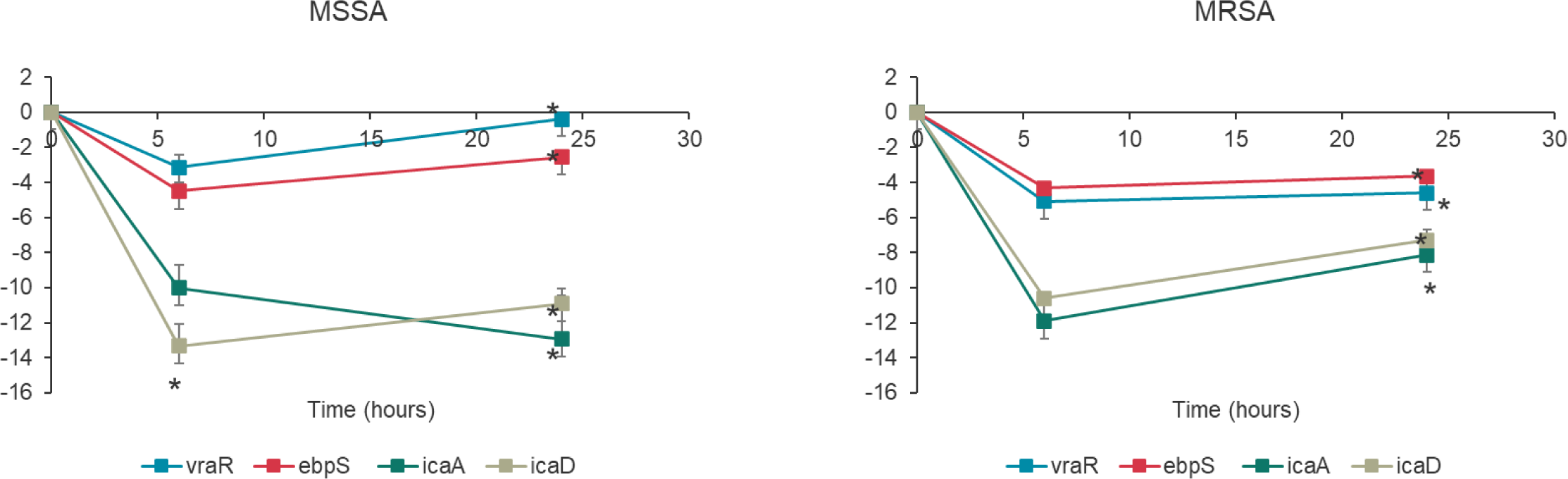
Regulation of stress responses, adhesion, and biofilm formation in implant colonizing MSSA and MRSA in vitro. Gene expression analysis of MSSA and MRSA biofilms adhered to the implant materials retrieved from the in vitro stainless-steel plates at time points indicated. Gene expression was normalized to 16srRNA expression. The gene expression was normalized to the planktonic *S. aureus* expression profiles. Error bars represent standard deviation (n=3). * Indicates p-value <0.05

## Discussion

Periprosthetic joint infection (PJI) remains a crucial clinical challenge that impacts implant longevity in patients, and results in high morbidity and mortality[37,38]. The persisting populations of bacterial biofilms colonizing the implant surface and surrounding tissue increase the severity of PJI and elevate the risk of recurring infections[39]. A comprehensive understanding of the biofilm establishment timeline, the evolution of drug resistance mechanisms, and the emergence of heterogenous phenotypes and genotypes using suitable in vivo and in vitro PJI models are key to devising effective eradication strategies that can curb bacterial resistance and persistence[18,40,41]. In this study, we investigated and characterized the pathogenesis and biofilm dynamics of a low-risk and high-risk infection in PJI. We aimed to longitudinally evaluate the risk of infection by employing in-vivo and in-vitro models. Our goal in vivo was to characterize the bacteria more extensively by incorporating both longitudinal analysis as well as the determination of resistance evolution. We aimed to elucidate the optimum therapy and therapeutic window to maximize the eradication of the bacteria based on the severity of PJI, simulated by using two strains with varying resistance. Our goal in vitro was to understand the relevance of the bacterial dynamics to in-vivo bacterial behavior and to determine methods to increase this relevance.

Bacterial contamination of the prosthetic components often occurs during their implantation. Despite the wide acceptance of the seminal concept of ‘race for the surface’ between the bacteria and host, recent findings indicate that the severity of the resulting infection largely depends on the bacterial strain, bacterial load, and the environment[42–47]. In our study, the ATCC 12600 strain was selected to simulate an infection that is susceptible to commonly used antibiotics (low risk) in the clinic. L1101, a clinically isolated Mu50 strain with known resistance to penicillin, aminoglycosides, and vancomycin, was used to simulate a ‘high-risk’ infection. To establish sustainable infections in the rat model in vivo, a high bacterial inoculum (10^8^ CFU) was used[48], and an in-vitro implant infection (10^5^ CFU) was established to characterize bacterial dynamics on surfaces. In the subcutaneous model, preferential colonization of the peri-implant tissue was observed over the entire study period compared to consistently poor colonization of the subcutaneous plate implant despite seeding with high bacterial density. Under SEM, the biofilms of MSSA were sparse compared to MRSA, which validates the advantage of inherent resistance in evading host immune responses[49]. On the other hand, in the joint infection model, almost equally high colonization was observed for MSSA and MRSA on the screw implants as well as the surrounding tibial and femoral tissues until POD 7. These observations indicate a strong influence of the internal environment within the infected site in driving the biofilm growth dynamics and in sustaining microbial viability[50]. Moreover, the microbial viability on contaminated implanted screws was ∼3 logs higher than the subcutaneous implant, which was corroborated by the SEM observations. The increased SA colonization on screw implants further underlined the flexibility of the facultative anaerobe SA in thriving in hypoxic environments[51,52]. The biofilm growth dynamics observed on in-vitro screw and plate models were largely comparable to their respective counterparts in the in-vivo models despite a lower starting inoculum. The vital role of different factors such as the implant location, site of bacterial colonization (implant vs tissue), the phenotypic and genotypic differences between bacteria, and the presence of vascularization in influencing biofilm dynamics in vivo is emphasized. The observations also facilitated the comparison between in-vivo and in-vitro infection and underscored the need for more diverse models to capture the biofilm growth dynamics and strain-specific infection outcomes.

Timely determination of the identity and antibiotic susceptibility profile of contaminating organisms can be crucial in making clinical dosing decisions to combat PJI[53–55]. Usually, the minimum inhibitory concentration (MIC) of infecting organisms directs the empirical antibiotic treatment regimens[13,56]. This approach, although widely accepted, relies on the assumption that the infecting organisms exist in a ‘planktonic state.’ In reality, microbes form complex biofilms during pathogenesis, wherein their physiology, drug susceptibility, and response to the host environment are continuously evolving, resulting in phenotypic and genotypic heterogeneity[22,57,58]. Our study highlights the profound impact of biofilm formation on the antibiotic susceptibility of MSSA and MRSA in periprosthetic joint infection. In the subcutaneous infection model, the tissue colonizing MSSA rapidly acquired gentamicin resistance as early as POD 1 (50 × MIC) and by POD7 required almost 100 × MIC to observe >3log reduction. This provided a critical insight regarding the minimal antibiotic dose for bacterial eradication, even for a low-risk susceptible strain of infecting bacteria. In comparison, the inherently resistant MRSA expectedly remained highly resistant to gentamicin (>500 × MIC) until POD 7. Notably, the decrease in the bacterial viability and an increase in the susceptibility of the tissue-colonizing MRSA on POD21 strongly suggests that the host response can be effective against a high-risk organism and the timeline of the infection is a vital component in infection characterization. For the implant-adhered bacteria, the MSSA adhered to the implant, gradually acquired resistance, and maintained viability over time, whereas the MRSA maintained its resistance profile throughout the study period. This behavior contrasts with the bacteria colonizing the tissue where there is more variation in growth dynamics and susceptibility profiles of the bacterial population, presumably due to an increased interaction with the host.

In the joint infection model, MSSA’s susceptibility to gentamicin evolved to >10 × MIC (implant-adhered) and 50 × MIC (tissue-colonized) within 7 days of infection which was comparable to previously published findings[59]. The high-risk MRSA showed consistently high resistance to the antibiotic over the entire study period. Compared to the subcutaneous infection model, the antibiotic susceptibility profile of MSSA in the joint infection model was not drastically elevated, which further emphasized the model-based longitudinal differences in infection risk stratification. In contrast to the MSSA populations in vivo, implant adhered MSSA acquired rapid and higher resistance to gentamicin within 24 hours (500 × MIC) in vitro. The data from plate-adhering MSSA and MRSA were comparable to the bacterial profiles observed until POD 3 and POD 7, respectively, in the subcutaneous infection model despite the absence of host factors in the in-vitro model. The in vitro infection study simulated more advantageous conditions for the bacteria in an implant-associated infection without host immunity. Based on the antibiotic susceptibility data from the three different infection models, a high initial dosing of antibiotics (>100 × MIC) is required during the early stages of infection that should be sustained for a prolonged period irrespective of the susceptibility profiles of the bacterial strain. Antibiotic-loaded implant materials that have been developed to locally release high antibiotic concentrations in a sustained manner could be best suited for this application[60–62]. For inherently resistant infections, it is advisable to use stronger tools such as the synergistic use of antibiotic and non-antibiotic compounds to enhance the antibacterial activity as early as possible to achieve effective eradication[36,63,64]. The study highlights the importance of diagnostics for characterizing the infecting bacteria *in situ* and developing more nuanced situation-specific and environment-specific guidelines for antibacterial treatment.

Current PJI diagnosis is largely limited in the identification and determination of antibiotic susceptibility of the causative organism[34]. The prophylactic and treatment models rely on the physiological attributes that have been determined using data largely from in-vitro studies[12]. The molecular status of bacterial populations within biofilms undergoes strain and maturity-dependent changes in their stress responses, cell wall constituents, metabolism, slime, and adhesin production, resistance to antibiotics, and immune responses[20,65,66]. Besides, the bacterial populations differ in their physiology depending on the site of colonization[67]. This adds a layer of complexity in the prevention and treatment approaches required. Taken together, the ‘status’ of the infecting bacterial strains (with inherent and acquired antibiotic resistance) and the site of colonization could directly correlate with the risk and severity of the infection[31,68]. Thus, there is a lot of uncertainty in predicting the treatment outcomes in specific cases of infections. To address this knowledge gap, the molecular signatures of the MSSA and MRSA bacterial populations colonizing implants in vivo, and in vitro were characterized. In the subcutaneous model, the stress response associated with maintaining bacterial cell wall integrity was triggered early on for tissue colonized MSSA (POD1) when compared to MRSA (POD21) supporting the idea that there is a lack of alternative strategies in low-risk strain and the multiplexed mechanisms present in high-risk strain for evading host-mediated targeting[69–71]. In contrast, both strains showed consistent upregulation of adhesin expression over time which revealed that bacterial populations actively increased adhesion on the tissue. The biofilm formation was more pronounced in MRSA compared to MSSA which is associated with high biofilm-forming properties attributed to resistant strains of *S. aureus*[72]. This finding significantly correlated to the bacteria counts in the tissue and to the SEM observations of MRSA on SS plates. For implant-adhered bacteria, due to the limited RNA availability, we were able to capture only bacterial *16srRNA* expression levels which were supportive of strain-specific biofilm viability from bacterial numbers and SEM images.

In the joint infection model, the bacterial stress response to the in-vivo environment was also found to be strain-dependent with the upregulation of the *vraR* gene for both MSSA and MRSA within 7 days. Strikingly, the adhesin gene expression was only consistently upregulated for MSSA and not for MRSA, which indicated the possibility of an alternative site-dependent mechanism in MRSA mediating tissue colonization[73]. In contrast to the subcutaneous infection model, the biofilm-associated *ica* genes were all upregulated throughout the study period with strain-specific longitudinal differences. The biofilm gene expression data validated the bacterial viability and SEM observations of the SS screw implant indicating the unique site-specific advantage facilitating increased biofilm survival and immune cell evasion[30,74]. Similar to the subcutaneous infection model, due to the limitation of biofilm RNA from the screw implants, only the presence of *16srRNA* could be validated. For the in-vitro infection model, we were able to pool multiple implant materials colonized by bacteria to assess their molecular status. Due to the absence of any in-vivo or environmental factors, the implant material-adhered bacterial expression of *vraR* and *ebpS* was comparable to the planktonic bacteria. The gene expression levels of biofilm-associated genes were significantly downregulated compared to planktonic bacteria indicating the limitation of in-vitro models without the integrated response to the immune system to capture and simulate the biofilm-associated changes in bacteria colonizing the implant[67,75].

Our in-vivo studies were limited by the number of implants. Despite implanting 6 implants per animal in the subcutaneous model, the determination of MBEC and gene expression of the bacteria on the implants were still limited by the low bacteria count. In addition, we only had one implant per animal to work within the joint infection model, further limiting our analysis. Our in-vitro studies were also limited to growth in one type of medium, which is likely to be a strong determinant of bacterial dynamics and evolution of resistance[76,77].

## Conclusion

The role of inherent and biofilm maturity-associated antibiotic resistance, and site-specific resistance profiles of *S. aureus* in determining the risk and severity progression of PJI was captured using three different infection models. This could aid in determining a suitable ‘therapeutic window’ for clinical dosing guidelines when encountering a low-resistance or high-resistance infection. Our study has also provided crucial insights into evaluating the translational value of in-vivo and in-vitro PJI models. The in-vivo infection studies using two different models revealed vital information on the strain-specific bacterial colonization patterns, evolution of resistance, and physiological differences that are governed by the site of infection. Immune response markers correlating with the infection status could serve as an additional resource to determine the effective concentrations of antibiotic required. Our results suggest implantation at the desired site is required for relevant efficacy testing of anti-infective materials. Even though in-vitro implant infection models are important and practical for studying biofilm dynamics on materials, the absence of in-vivo factors limits their translational value.

## EXPERIMENTAL SECTION

### 1. Bacteria preparation

Gentamicin-susceptible *S. aureus* ATCC 12600 (MSSA) and Gentamicin-resistant *S. aureus* L1101 (MRSA) were used in this study (Table 1, supplementary file 1). The bacterial glycerol stocks at −80°C were grown in tryptic soy agar plates (TSA) for 18-24 hours at 35°C to achieve optimum growth. *S. aureus* colonies were inoculated in tryptic soy broth (TSB) and cultured overnight to obtain 10^9^ CFU/mL. The bacterial broth cultures were subjected to 10,000 × g centrifugation and pellets were resuspended in sterile PBS. The bacterial suspension in PBS was further diluted to 10^8^ CFU in sterile PBS and 10^5^ CFU/mL in sterile TSB before all animal infection experiments and in vitro experiments, respectively.

### 2. Animal study

#### 2.1 Ethics statement

The animal study design and protocols were approved by the Institutional Care and Use Committee of Massachusetts General Hospital (2021N000127).

#### 2.2 Subcutaneous infection model

Male Sprague-Dawley rats (Charles River, Wilmington, MA) weighing 350-400g were randomly divided into two groups. 316L stainless steel plates (10 × 3 × 1mm; n=6 in each animal) were subcutaneously implanted on each rat dorsum and 10^8^ CFU of gentamicin-sensitive MSSA (n=34 rats), gentamicin-resistant MRSA (n=33) were inoculated into each of the 6 subcutaneous pockets. Non-infected group (n=14) served as a control for the study. All rats were given facility chow and water ad libitum. Rats were anesthetized with 1-3% isoflurane in 1 L of O2/min, and 0.05 mg/kg IP buprenorphine was administered 30 minutes before surgery. All groups were sacrificed on postoperative days (POD) 1, 3, 7, and 21 (Table 2, Supplementary File 1)

### 2.3 Joint infection model

Male Sprague Dawley Rats were randomly assigned to ‘low-risk’ gentamicin-sensitive MSSA, ‘high-risk’ gentamicin-resistant MRSA infection groups (n=3/day) and control group (n=1/day). 10^8^ CFU of bacteria were inoculated into the intercondylar canal drilled in the tibia after which, a stainless-steel screw (1.3 mm diameter, 8 mm in length) was implanted into the contaminated canal. All rats were given facility chow and water ad libitum. Rats were anesthetized with 1-3% isoflurane in 1 L of O2/min, and 0.05 mg/kg IP buprenorphine was administered 30 minutes before surgery. The animals were sacrificed on POD 1, 3, and 7 (Table 3, supplementary file 1).

### 3. Ex-vivo and in-vitro determination of bacterial colonization

The stainless-steel plates and screws (SS) were retrieved from the animals at specific time points (POD 1, 3, 7, 21, and POD 1,3,7 respectively) and the explants were washed using sterile 1× phosphate-buffered saline (PBS) to remove any non-adherent bacteria and debris. The explants were then transferred to 1.5 mL tubes and subjected to sonication for 40 minutes in 1 mL PBS to dislodge the adherent bacteria. The sonicate was then diluted and plated on tryptic soy agar plates and incubated for 18-24 hours at 35°C. The adherent bacteria count was determined the following day using the colony counting method. The peri-implant tissue surrounding the implant was harvested at the same time points and homogenized using a Tissue Disruptor (TissueRuptor, Qiagen). The homogenate was diluted and plated on tryptic soy agar plates and incubated for 18-24 hours at 35°C. The tissue-colonized bacteria were determined the following day using the colony count method.

To determine in-vitro biofilm dynamics, staphylococcal bacterial suspension [10^5^ CFU/mL] in 1 mL of Luria-Bertani (LB) broth was inoculated on 316 Stainless Steel (SS) plates [10 × 3 × 1 mm] or screws placed within 24-well plates. The materials were statically incubated for an indicated period (6 and 24 hours) at 35°C. At each time point, the spent media was removed, and the materials were washed thrice using sterile 1× PBS. The surfaces were transferred to 1.5 mL tubes and subjected to sonication for 40 minutes in 1 mL PBS to dislodge the adherent bacteria, The sonicate was then plated on tryptic soy agar plates and incubated for 18-24 hours at 35°C. The adherent bacteria count was determined the following day by the colony counting method.

### 4. Ex-vivo and in-vitro determination of minimum biofilm eradication concentration

The stainless-steel plates and screws (SS) were retrieved from the animals at specific time points (POD 1, 3, 7, 21 and POD 1, 3, 7 respectively) and the plates were washed using sterile 1× PBS to remove any non-adherent bacteria and debris. The explants were then exposed to a range of gentamicin concentrations in 10% LB [1, 10, 50, 100, 200, and 500 µg/mL for SS plates; 10 (MSSA) and 300 µg/mL (MRSA) for screws)] for 24 hours at 35°C. The explants were then washed using sterile 1× PBS and subjected to sonication for 40 minutes in 1 mL PBS to dislodge the adherent bacteria. The sonicate was then diluted and plated on tryptic soy agar plates and incubated for 18-24 hours at 35°C. The adherent bacteria count was determined the following day using the colony counting method. The retrieved peri-implant tissue was rinsed once with sterile PBS and exposed to a range of gentamicin concentrations in 10% LB [1, 10, 50, 100, 200, and 500 µg/mL (subcutaneous infection study); 10, 50, 100, 300 and 500 µg/mL (joint-infection study)] for 24 hours at 35°C. The tissues were washed with sterile PBS and were sonicated for 40 minutes and plated. The bacterial viability was determined the following day using the colony count method.

To determine MBEC of in vitro-formed biofilms, staphylococcal biofilms were grown for a period of 6 and 24 hours on plates and screws as previously described (section 3). The spent media was removed at each timepoint respectively and the materials were washed thrice with PBS to remove all non-adherent bacteria. The surfaces were then placed in a fresh 24-well plate containing a range of gentamicin concentrations [0.5, 1, 5, 10, 20, 40, 60, 80, 100, 200, 300, 400, 500 µg/mL (SS plates); 1, 5, 10, 50, 100, 300, 500 (SS screws)]. Further to drug exposure for 24 hours at 35°C, the surfaces were gently rinsed thrice using PBS and were transferred to 1.5ml tubes containing 1ml PBS. The materials underwent sonication for 40 minutes and the adherent bacteria count was determined using the spread plate method. MBEC was determined as the concentration that achieved >3log10 reduction in adherent bacteria count.

The growth frequency observed from each replicate for each concentration tested was calculated as %growth frequency and heatmaps were generated using the Complex Heatmaps package in R studio.

### 5. Gene expression analysis

The peri-implant tissue retrieved from each time point was subjected to a modified Qiagen RNeasy extraction protocol to extract bacteria RNA. Briefly, 15-30mg tissue samples were homogenized and subjected to lysis using RLT buffer. The lysate was transferred to a tube and was further subjected to enzymatic and mechanical lysis using lysostaphin (200 µg/mL), proteinase K, and acid-washed beads. RNA extraction was performed using the RNeasy spin column method according to the manufacturer’s instructions. To determine in vitro and in vivo adherent bacteria gene expression the bacteria were harvested from SS plates (n=10 for in vitro model; n=6 from in vivo model) and screws (n=4 from in vivo model) for each condition by subjecting the materials to 40 min sonication. The sonicate fluid was pooled and pelleted by centrifuging at 10,000 ×g for 10 mins. The pellet was then subjected to mechanical and enzymatic lysis and the total RNA was extracted using RNeasy Power Biofilm RNA extraction kit for gram-positive bacteria. The samples were subjected to real-time quantitative PCR for *icaA*, *icaD*, *ebpS*, and *vraR* genes for *S. aureus* using specific primers listed (Table 4, supplementary file 1). The Cq values were normalized to *S. aureus* 16srRNA expression. Gene expression analysis of tissue-colonizing and implant-adhered bacteria relative to planktonic bacterial expression was performed using the 2^(-ΔΔCt)^ method.

### 6. Scanning Electron Microscopy

Scanning electron microscopy was performed on the implant materials retrieved from rats on POD3 and from in-vitro experiments[36]. The implant materials with adherent bacteria were fixed using 2.5% glutaraldehyde in 0.1M PBS for 48 hours. The plates were then washed twice for 10 mins with PBS. The adherent bacteria were then treated with osmium tetroxide (OsO^4^) 2% + Ruthenium red 0.2% 1:1 solution for a period of 1hr. The samples were washed twice thoroughly with distilled water for 10 mins. Further to this the samples were treated with 1% Tannic acid for 30 mins and then washed twice with distilled water for 10 mins each. The prepared samples were imaged at 15-20 kV, high vacuum (Zeiss FESEM Ultra Plus).

### 7. Statistical analysis

The gene expression studies were performed in triplicates and the dataset was analyzed using Student’s T-Test (paired). The p-value was calculated and the lowest significant score of 0.05 was considered statistically significant.

## Author contributions

AS, EO, and YF conceptualized and conducted the experimental design for the study. AS, PT, MM, PJ, FM, and DK performed the data acquisition and SEM visualization. YF, JY, and SL performed the animal surgery protocols. KKW and NI performed SEM visualization. AS analyzed and interpreted the data. AS, EO, and OKM wrote, reviewed, and edited the manuscript. All authors read and approved the final manuscript.

## Acknowledgments

This study was funded by National Institutes of Health Grant R01AR077023. The funder played no role in the study design, data collection, analysis, and interpretation of data, or the writing of this manuscript. The authors thank Jean Yuh and Sashank Lekkala for their technical assistance during animal surgery and Dr. Kerry Laplante at the University of Rhode Island for providing the clinical MRSA strain. This work was performed in part at the Harvard University Center for Nanoscale Systems (CNS); a member of the National Nanotechnology Coordinated Infrastructure Network (NNCI), which is supported by the National Science Foundation under NSF award no. 651 ECCS-2025158.

## Data availability

All data generated or analyzed during this study are included in this published article [and its supplementary information files]

