## Supplementary File for "Investigating the translational value of Periprosthetic Joint Infection (PJI) models to determine the risk and severity of Staphylococcal biofilms"

| Table 1: Antibiotic susceptibility data of bacterial strains used in the study | | | |
|---|---|---|---|
| **Bacterial strain** | **Methicillin** | **Gentamicin** | **Vancomycin** |
| ATCC 12600 | Sensitive | Sensitive | Sensitive |
| L1101 (Mu50) | Resistant | Resistant | Resistant (Intermediate) |

| Table 2: Number of animals in each study group and samples retrieved for Subcutaneous infection model | | | | | | | |
|---|---|---|---|---|---|---|---|
| | **Plate culture** | **Tissue culture** | **Plate MBEC** | **Tissue MBEC** | **Tissue gene expression** | **Plate SEM** | **Plate gene expression** |
| **POD1**<br>12600 N= 7<br>L1101 N=7<br>NIC N=3 | 12600 N=4<br>L1101 N=4<br>NIC N=18 | 12600 N=15<br>L1101 N=15<br>NIC N=3 | 12600 N=2 per concentration tested<br>L1101 N=2 per conc tested<br>*2 independent experiments | 12600 N=2 per concentration tested<br>L1101 N=2 per conc tested<br>*2 independent experiments | 12600 N=3<br>L1101 N=3 | | |
| **POD3**<br>12600 N= 9<br>L1101 N=9<br>NIC N=5 | 12600 N=4<br>L1101 N=4<br>NIC N=18 | 12600 N=15<br>L1101 N=15<br>NIC N=3 | 12600 N=2 per concentration tested<br>L1101 N=2 per conc tested<br>*2 independent experiments | 12600 N=2 per concentration tested<br>L1101 N=2 per conc tested<br>*2 independent experiments | 12600 N=3<br>L1101 N=3 | 12600 N=2<br>L1101 N=2 | 12600 N=2<br>L1101 N=2<br>6 plates /group |
| **POD7**<br>12600 N= 7<br>L1101 N=7<br>NIC N=3 | 12600 N=4<br>L1101 N=4<br>NIC N=18 | 12600 N=15<br>L1101 N=15<br>NIC N=3 | 12600 N=2 per concentration tested<br>L1101 N=2 per conc tested<br>*2 independent experiments | 12600 N=2 per concentration tested<br>L1101 N=2 per conc tested<br>*2 independent experiments | 12600 N=3<br>L1101 N=3 | | |
| **POD21**<br>12600 N= 11<br>L1101 N=10<br>NIC N=3 | 12600 N=14<br>L1101 N=8<br>NIC N=18 | 12600 N=11<br>L1101 N=10<br>NIC N=3 | 12600 N=2 per concentration tested<br>L1101 N=2 per conc tested | 12600 N=2 per concentration tested<br>L1101 N=2 per conc tested | 12600 N=11<br>L1101 N=10 | | |

| Table 3: Number of animals in each study group and samples retrieved for Joint infection model | | | | | | | |
|---|---|---|---|---|---|---|---|
| | **Screw culture** | **Tissue culture** | **Screw MBEC** | **Tissue MBEC** | **Tissue gene expression** | **Screw SEM** | **Screw gene expression** |
| **POD1**<br>12600 N= 3<br>L1101 N=3<br>NIC N=1 | 12600 N=1<br>L1101 N=1<br>NIC N=1 | Femoral tissue and Tibial tissue<br><br>12600 N=3<br>L1101 N=3<br>NIC N=2 | 12600 N=1 per concentration tested<br>L1101 N=1per concentration tested | 12600 N=1 per concentration<br>L1101 N=1 per concentration<br>Total N per infection group =3 | 12600 N=3<br>L1101 N=3 | | |
| **POD3**<br>12600 N= 5<br>L1101 N=3<br>NIC N=1 | 12600 N=1<br>L1101 N=1<br>NIC N=1 | Femoral tissue and Tibial tissue<br><br>12600 N=3<br>L1101 N=3<br>NIC N=2 | 12600 N=1 per concentration tested<br>L1101 N=1per concentration tested | 12600 N=1per concentration<br>L1101 N=1 per concentration<br>Total N per infection group =3 | 12600 N=3<br>L1101 N=3 | 12600 N=2 | 12600 N=4 |
| **POD7**<br>12600 N= 3<br>L1101 N=3<br>NIC N=1 | 12600 N=1<br>L1101 N=1<br>NIC N=1 | Femoral tissue and Tibial tissue<br><br>12600 N=3<br>L1101 N=3<br>NIC N=2 | 12600 N=1 per concentration tested<br>L1101 N=1per concentration tested | 12600 N=1per concentration<br>L1101 N=1 per concentration<br>Total N per infection group =3 | 12600 N=3<br>L1101 N=3 | | |

| Table 4: List of primers used in the study | |
|---|---|
| *vraR* | FP 5’-AACTCTGCGCGCTTTTTCAT-3’ |
| | RP 5’-ATATCGCCGATGCAGTTCGT-3’ |
| *icaA* | FP 5’-TTGTCGACGTTGGCTACTGG-3’ |
| | RP 5’-GCGTTGCTTCCAAAGACCTC-3’ |
| *icaD* | FP 5’-CGCTATATCGTGTGTCTTTTGGA-3’ |
| | RP 5’-TCGCGAAAATGCCCATAGTT-3’ |
| *ebpS* | FP 5’-TACTTTGGCCATGCCACCTT-3’ |
| | RP 5’-TGCTTCTGCCGCTTCAAAAC-3’ |
| *16srRNA* | FP 5’-AGACCAGAAAGTCGCCTTCG-3’ |
| | RP 5’-TCAACCGTGGAGGGTCATTG-3’ |

Table 5: Cq values obtained for implant-associated bacterial gene expression

| Sample | Infection group | Target gene | Cq value |
|---|---|---|---|
| Joint infection model<br>Screw- adherent bacteria (POD3) | MSSA | *16srRNA* | 29.67 |
| | | *16srRNA* | 29.92 |
| | | *16srRNA* | 31.38 |
| | | *icaA* | N/A |
| | | *icaA* | N/A |
| | | *icaA* | N/A |
| | | *icaD* | N/A |
| | | *icaD* | N/A |
| | | *icaD* | N/A |
| | | *ebpS* | N/A |
| | | *ebpS* | N/A |
| | | *ebpS* | N/A |
| Subcutaneous infection model<br>Plate-adherent bacteria (POD3) | MSSA | *16srRNA* | 34.57 |
| | | *16srRNA* | 37.59 |
| | | *16srRNA* | 38.86 |
| | | *icaA* | N/A |
| | | *icaA* | N/A |
| | | *icaA* | N/A |
| | | *icaD* | N/A |
| | | *icaD* | N/A |
| | | *icaD* | N/A |
| | | *ebpS* | N/A |
| | | *ebpS* | N/A |
| | | *ebpS* | N/A |

| | | | |
|---|---|---|---|
| Subcutaneous infection model Plate-adherent bacteria (POD3) | MRSA | *16srRNA* | 21.78 |
| | | *16srRNA* | 20.02 |
| | | *16srRNA* | 22.49 |
| | | *icaA* | N/A |
| | | *icaA* | N/A |
| | | *icaA* | N/A |
| | | *icaD* | N/A |
| | | *icaD* | N/A |
| | | *icaD* | N/A |
| | | *ebpS* | N/A |
| | | *ebpS* | N/A |
| | | *ebpS* | N/A |
